# Invasive freshwater snails form novel microbial relationships

**DOI:** 10.1101/2020.04.29.069013

**Authors:** L. Bankers, D. Dahan, M. Neiman, C. Adrian-Tucci, C. Frost, G.D.D. Hurst, K.C. King

**Affiliations:** University of Iowa, Department of Biology, Iowa City, IA, USA; Stanford University, School of Medicine, Stanford, CA, USA; University of Liverpool, Institute of Integrative Biology, Liverpool, UK; University of Oxford, Department of Zoology, Oxford, UK

**Author notes:** **Corresponding author:** Laura Bankers. LB and DD contributed equally to this work and should be considered joint first authors. University of Colorado Anschutz Medical Campus, Aurora, CO, USA. 1150 4^th^ St. SW Apt. 607 Washington DC, USA.

**Keywords:** *Potamopyrgus antipodarum*, invasion, microbiome, coevolution, symbiosis, infection

## Abstract

Resident microbes (microbiota) can shape host organismal function and adaptation in the face of environmental change. Invasion of new habitats exposes hosts to novel selection pressures, but little is known about the impact of invasion on microbiota and the host-microbiome relationship after this transition (e.g., how rapidly symbioses are formed, whether microbes influence invasion success). We used high-throughput 16S rRNA sequencing of New Zealand (native) and European (invasive) populations of the freshwater snail *Potamopyrgus antipodarum* and found that while invaders do carry over some core microbial taxa from New Zealand, most of their microbial community is distinct. This finding highlights that invasions can result in the formation of novel symbioses. We further show that the native microbiome is composed of fewer core microbes than the microbiome of invasive snails, suggesting that the microbiota is streamlined to essential members. Together, our findings demonstrate that microbiota comparisons across native and invasive populations can reveal the impact of a long coevolutionary history and specialization of microbes in the native host range, as well as new associations occurring after invasion. We lay essential groundwork for understanding how microbial relationships affect invasion success and how microbes may be utilized in the control of invasive hosts.

## INTRODUCTION

All organisms are home to numerous microbes. The relationship between hosts and their microbial communities (microbiota) can play a role in key host functions (1, 2) from nutrition (3) to behaviour (4) and immune responses (5). The microbiota has also been shown to impact host evolution (6, 7). Microbial community composition can be determined by host traits such as genotype (8), sex (9), infection of the host by parasites or pathogens (10), environmental conditions (6, 11), and genotype by environment interactions (12). Therefore, both host organisms and the microbes on which they depend for vital processes can both be impacted by environmental changes, with potential consequences for host adaptation (13).

When species invade novel habitats, they are exposed to new environmental conditions and selection pressures. Because host and microbe fitness can be mutually dependent, invasion is predicted to influence both the host and its microbiome (6). The fitness effects of microbes on their host may affect invasion success and the potential for subsequent adaptation to a new habitat (14). The performance of introduced species could also be altered by the availability of microbes outside their native range if there are core microbes that the host needs to thrive. If host-associated microbes are not all carried over from the native environment, lost upon invasion, and/or unavailable in the new environment, the diversity of the microbiomes of invasive lineages could be low. The invader’s microbiome might thus only represent a subset of that residing in the native lineages (15, 16). This pattern is directly analogous to the loss of genetic diversity that often characterizes invasive lineages (15, 17). Whether such subsampling occurs at the microbiome level remains an important open question with implications for host fitness in the new environment.

New invasions may also affect the level of specialization of microbiota relative to hosts in the native range. This outcome could be a consequence of the comparatively short time period for coevolution between the invasive host and newly acquired microbes vs. the microbes from the native environment (15, 16). These effects of colonization could influence microbiome structure. These consequences could be harmful for the host, such as in a situation where suitable mutualistic microbes are not in the new habitat. It is also possible that invasive hosts form new symbioses with the microbes available (18). A more diverse microbiota may result if there is decreased immune-mediated control of novel microbes (19). By this logic, we might therefore expect native host individuals to have a less rich microbiota. In particular, established and stable microbial communities, in which novel competitive interactions between microbes have long since played out, are likely to be of lower diversity (20). This prediction might be particularly likely to hold if the host has evolved to cultivate beneficial associations (19). Addressing whether this prediction is met requires characterization of how invasion impacts microbial diversity and composition.

Molluscs are one of the most species-rich phyla (21). They represent an ecologically and economically important class of invaders (22, 23), however, little is known about the relationships between molluscs and their symbiotic microbes with the exception of a handful of recent studies (24, 25). Here, we leveraged a powerful model system for host-parasite coevolution, the New Zealand freshwater snail *Potamopyrgus antipodarum*, to characterize the impact of invasion on host microbiota. There is substantial across-lake population variation in microbiome composition among native *P*. *antipodarum* (26). *Potamopyrgus antipodarum* in these native New Zealand populations also feature wide individual-level variation in reproductive mode (sexual vs. asexual) and ploidy level (sexual snails are diploid while asexual snails are polyploid (triploid or tetraploid); (27, 28)), and are coevolving with sterilizing trematode parasites (29, 30). By contrast, invasive lineages of *P*. *antipodarum* in Europe and North America are polyploid asexuals with reduced genetic variation compared to the native range (17, 31) and have escaped from parasite infection (32, 33).

We conducted 16S rRNA sequencing of field-collected *P*. *antipodarum* from waterbodies across New Zealand and Europe to assess changes in microbiota diversity and composition across and within native and invasive host populations. We hypothesized that, 1) invasive *P*. *antipodarum* would have a subset of native microbial diversity because invaders are often subsamples of native diversity, which could also likely lead to lower microbial diversity (analogous to patterns often described for genetic variation post-invasion). Lower microbial diversity could arise because they simply have had less time in the new environment to acquire new symbioses; 2) key symbionts needed to thrive should be maintained in the new environment; 3) invaders will have fewer conserved microbes than native counterparts as a consequence of relatively recent contact with symbionts in the new environment, and therefore, relatively low time for the evolution of specialization. To our knowledge, our study of *P*. *antipodarum* is the first comparison of diversification in host-associated microbial communities between native and invasive molluscs. We also address associations between the fundamental and often-variable organismal traits of sex, reproductive mode, and infection status on microbiota within those host populations. More broadly, the results of this study provide an important starting point to evaluate whether and how microbes could be utilized in efforts to control invasive species.

### Study System

Native New Zealand *P*. *antipodarum* are primarily found in freshwater lakes and streams (34). These snails have gained prominence as a model for studying the evolutionary maintenance of sexual reproduction (27, 28) as well as host-parasite coevolution (27, 30). Multiple triploid and tetraploid asexual lineages have been separately derived from diploid sexual conspecifics (27, 28), and New Zealand populations are commonly infected by a sterilizing trematode parasite, *Atriophallophorus winterbourni* (formerly *Microphallus livelyi*) (27, 30, 35). A previous study hinted that the microbiome composition of these snails varies substantially among native populations and between sexuals and asexuals (26). However, these snails were either laboratory cultured or housed in a laboratory for several months before harvest (Maurine Neiman, personal communication) meaning that whether these results hold in snails sampled directly from the field remains unclear.

Only asexual lineages of *P*. *antipodarum* are highly successful invaders – why this is remains unclear. Nevertheless, *P*. *antipodarum* can survive a range of harsh conditions that may facilitate travel to and establish in new habitats (22, 36). Invaders are susceptible to few biological enemies (22), passing alive through the digestive systems of trout (37) and are only rarely infected by parasites (36). Once they invade, these snails can influence ecosystems due to rapid population growth (22, 38), high population densities (32, 39), and competitive exclusion of native invertebrates (22, 38). Invasive *P*. *antipodarum* can even harm the native predators that ingest them (37).

## MATERIALS AND METHODS

### Sample Collection, Processing, and Sequencing

We collected adult *Potamopyrgus antipodarum* during warm seasons from shallow (lake depth < 1 m) rocks and vegetation from three New Zealand collection sites (early January 2015) and ten European collection sites (late May 2016) (Table 1; Fig. S1). Snails were maintained in identical 15 L tanks separated by source population in a 16° C room with a 16:8 hour light:dark cycle for less than one month before dissection. Snails were fed dried *Spirulina* cyanobacteria *ad libitum* (40).

**Table 1.** Sample collection sites. The coordinates and location names of sample sites for both native and invasive snails are list below. Ploidy is a proxy for reproductive mode; diploids are sexual, polyploids are asexual. We also include the sex, infection status, and number of individuals sequenced from each condition and location and condition.

| Country | Location | Geographic Coordinates | Reproductive Mode | Infection | Sex | No. Snails |
| --- | --- | --- | --- | --- | --- | --- |
| New Zealand | Lake Alexandrina | 43° 56' 12.9" S, 170° 27' 35.4" E | Sexual | Infected | Female | 9 |
|  |  |  | Sexual | Infected | Male | 2 |
|  |  |  | Sexual | Uninfected | Female | 22 |
|  |  |  | Sexual | Uninfected | Male | 19 |
|  |  |  | Asexual | Unknown | Female | 16 |
|  |  |  | Asexual | Infected | Female | 10 |
|  |  |  | Asexual | Uninfected | Female | 20 |
| New Zealand | Lake Kaniere | 42° 48' 21.8" S, 171° 7' 40.5" E | Sexual | Infected | Female | 2 |
|  |  |  | Sexual | Uninfected | Female | 6 |
| New Zealand | Lake Rotoroa | 41° 47' 44.5" S, 172° 36' 22.4" E | Sexual | Infected | Female | 1 |
|  |  |  | Sexual | Uninfected | Female | 10 |
|  |  |  | Asexual | Infected | Female | 1 |
|  |  |  | Asexual | Uninfected | Female | 10 |
| Austria | Lake Mondsee | 47° 50' 37.7" N, 13° 20' 30.7" E | Asexual | Uninfected | Female | 17 |
| Belgium | Brakel | 50° 48' 12.1" N, 3° 45' 20.9" E | Asexual | Uninfected | Female | 12 |
| Belgium | Geraardsbergen | 50° 47' 30.4" N, 3° 54' 15.0" E | Asexual | Uninfected | Female | 13 |
| Belgium | Melle | 51° 1' 11.6" N, 3° 48' 59.6" E | Asexual | Uninfected | Female | 12 |
| Germany | Wrohe | 54° 16' 7.4" N, 9° 57' 35.9" E | Asexual | Uninfected | Female | 3 |
| Germany | Pulsen | 54° 19' 17.3" N, 10° 27' 9.8" E | Asexual | Uninfected | Female | 12 |
| Switzerland | Lake Geneva | 46° 30' 45.2" N, 6° 36' 27.4" E | Asexual | Uninfected | Female | 15 |
| United Kingdom | West Holme | 50° 40' 20.1" N, 2° 10' 29.03" W | Asexual | Uninfected | Female | 11 |
| United Kingdom | Bovington | 50° 41' 26.1" N, 2° 13' 29.02" W | Asexual | Uninfected | Female | 12 |
| United Kingdom | Oxford | 51° 45' 6.5" N, 1° 13' 0.35" W | Asexual | Uninfected | Female | 12 |

All snails were sexed and shells were removed prior to DNA extraction and assessed for *A*. *winterbourni* infection based on the presence of metacercariae cysts via dissection (41) (Table 1). Snails containing non-*A*. *winterbourni* infections were excluded from the study. All metacercariae were removed from infected snails using a micropipette. After dissection, we used flow cytometry (28, 42) to determine ploidy status as a proxy for reproductive mode of New Zealand samples (diploids are sexual and polyploids are asexual). Because it is already established that the European invasive lineages are virtually all polyploid asexuals (17, 43), we did not perform flow cytometry on these samples.

We used the Qiagen DNeasy Plant Mini Kit (QIAGEN Inc.) to extract DNA from whole snails (shells and metacercariae removed), following manufacturer protocol, but eluting DNA in 40 μl 100:1 TE buffer. The Plant kit was used as it better handles the polysaccharides present in snail mucus, compared to other DNA extraction kits. Samples with a 260/280 ratio > 1.6 and containing > 20 ng of total DNA, based on Nanodrop® 1000 (Thermo Fisher Scientific), were analysed on a 1% agarose gel. Samples that reached our quality criteria and produced clear gel bands were shipped to the W.M. Keck Center for Comparative Functional Genomics (University of Illinois at Urbana-Champaign) for sequencing. See Electronic Supplementary Methods for more details about sample collection and processing.

The 16S rRNA V4 region was amplified from the *P*. *antipodarum* microbiome gDNA using the 515F Golay-barcoded primers and 806R primers (44, 45). Samples were prepared in accordance with the standard Earth Microbiome Project 16S rRNA protocol (46). Please see our Electronic Supplementary Methods for PCR reaction mixtures and thermocycler conditions. gDNA was quantified using the Qubit 2.0 fluorometer (Thermofisher Scientific), amplicons were pooled at equimolar ratios (∼240 ng per sample), and amplicon pools were cleaned using the Qiagen PCR Purification Kit (QIAGEN Inc.). The multiplexed library was quality checked and sequenced with the MiSeq 2×250 bp PE v2 protocol at the W.M. Keck Center for Comparative Functional Genomics (University of Illinois).

### Computational and Statistical Analyses

We removed PhiX sequences from sequencing libraries using Bowtie2 (47) (PhiX genome obtained from: support.illumina.com/sequencing/sequencing_software/igenome.html). Demultiplexed paired-end fastq files were processed using DADA2 in R (3.4.0), with the suggested filtering and trimming parameters, as previously described (48). We then merged paired-end reads and constructed an amplicon sequence variant (ASV) table. We used the native implementation of the DADA2 Ribosomal Database Project (RDP) naïve Bayesian classifier (49) trained against the GreenGenes 13.8 release reference fasta (https://zenodo.org/record/158955#.WQsM81Pyu2w) to classify ASVs taxonomically.

We used phyloseq v. 1.16.2 to perform estimate_richness and vegan’s pd function to calculate alpha diversity measurements of observed ASVs, Shannon’s index, PD Whole Tree, Pielou’s evenness, and Chao1 (50), and to perform ordinations using PCoA on unweighted and weighted UniFrac distance scores (51). We used the Songbird reference frames approach to calculate taxa differentials (52). Data visualization and statistical analyses were performed in R; see Electronic Supplemental Methods for details.

We controlled for effects of snail sex, reproductive mode, and infection status (Table 1) when testing for geographic associations with microbiota by only conducting analyses on female uninfected asexual snails, allowing us to directly compare native and invasive snails without the confounding factors of sex, reproductive mode, and infection (comparisons of sex, reproductive mode, and infection status are described below). For alpha diversity analyses, we rarefied samples to 10,000 ASVs per sample and discarded two samples that had fewer reads than this threshold. To test covariate effects on microbiota alpha diversity, we used Welch’s Two Sample t-tests and adjusted *p*-values (“adj-*p*”) with a Bonferroni correction for multiple tests. We conducted beta diversity analyses on all ASVs after removing singletons. We normalized ASV counts by adding one and then loge-transforming ASV counts (48). To evaluate beta diversity we performed PCoA on the distance matrices built on the unweighted and weighted UniFrac scores of each sample (51).

We used Analysis of Similarity (ANOSIM; 999 permutations, R-statistics (R^2^) and exact *p*-values reported in Results) to evaluate effects of geographic location (sample site) on microbiota beta diversity. Because all European snails were female uninfected asexual snails and many New Zealand snails were sexual, infected, and/or male, we only performed the ANOSIM comparing Europe and New Zealand on female uninfected asexual snails.

To analyse population specificity, we focused on the core microbiome of snails within Europe or New Zealand, defined as taxa found in at least 90% of samples within Europe or New Zealand. We used a t-test to compare the proportion of reads that mapped as core microbiome taxa between the combined Europe and combined New Zealand sample groupings.

We performed a Permutational Multivariate Analysis of Variance Using Distance Matrices analysis (ADONIS) to test the effects of reproductive mode, sex, and infection status on microbiota beta diversity. The ADONIS analysis was limited to the Lake Alexandrina (New Zealand) sample site as it was the only site for which we were able to obtain all conditions (sexual, asexual, male, female, infected, and uninfected). ADONIS tests were conducted with 999 permutations.

We employed the Analysis of Composition of Microbes (ANCOM) algorithm to conduct a differential count analysis (53). These analyses were limited to the Lake Alexandrina (New Zealand) sample site for the reasons noted above. We tested the effect of sex in uninfected male vs. uninfected female sexual snails, tested the effect of reproductive mode on uninfected female sexual vs. uninfected female asexual snails, and tested the effect of infection status on asexual female trematode-infected vs. asexual female uninfected snails. We used machine learning to model microbiome classification by geography using female uninfected asexual snails and ASVs agglomerated phylogenetically using default settings (phyloseq tax_glom; h = 0.2). Machine Learning was trained on a random forest model using the caret package (v6.0-81) with a test split of 80:20 and fit a random forest classifier over the tuning parameter of snail origin (New Zealand or Europe). Please see the Electronic Supplemental Methods for more details on bioinformatic and statistical analyses.

## RESULTS

Variation in *P*. *antipodarum* microbiota was in large part driven by whether a snail was from Europe or New Zealand (Fig. 1; ANOSIM; R = 0.97; *p* < 0.01; Permutations: 999; ADONIS; R^2^ = 0.33, *p* < 0.01; weighted UniFrac dissimilarity distances are presented in Fig. S2). To validate this analysis, we performed a Mantel test on coordinate and microbiota distance matrices and again found geographic distance was significantly associated with microbiome distance (Mantel test; Observation = 0.13; *p* < 0.01). Within the two European and New Zealand regions, we did not find significant differences in microbial variation among sample sites. We thus attribute the geographic distance-to-microbiome distance correlation to major differences between microbiomes from the most geographically distant regions of Europe and New Zealand, rather than differences among sites within the two larger regions. To confirm that beta diversity differences between microbiomes of snails from Europe and New Zealand were not due to uneven sampling (10 European sample sites vs 3 New Zealand samples sites), we permuted PCoA on the distance matrices built on the unweighted UniFrac on all combinations of three Europe sites vs. the three New Zealand sites (Fig S3).

**Fig 1.**
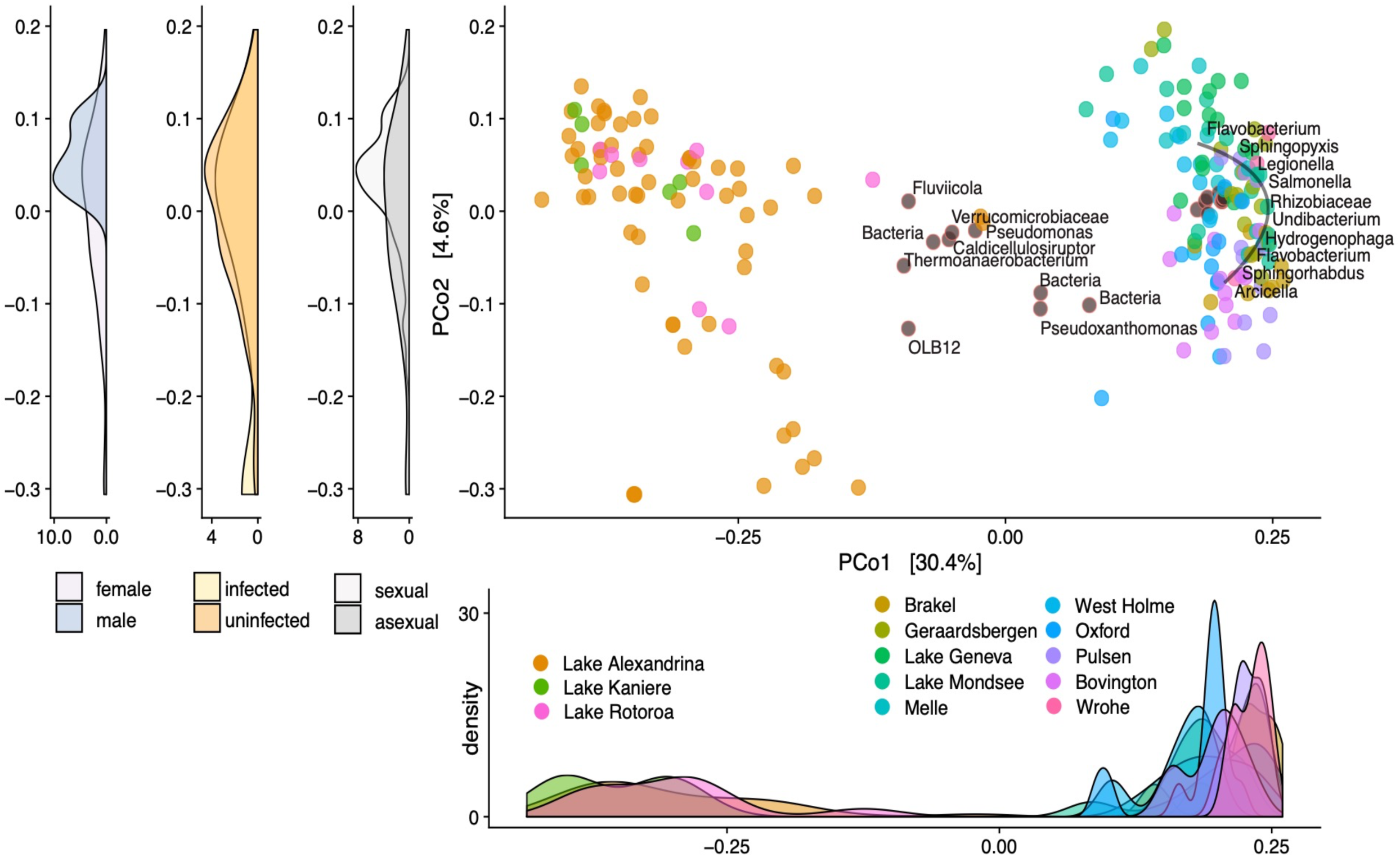
Snail microbiota ecosystem clustering. Biplot PCoA on unweighted UniFrac dissimilarity distances among snail microbiota profiles across European and New Zealand sample sites. Each point represents a single snail and points are colour coded based on whether a point is a microbial taxon (grey filled with outline colour indicating sample site) or snail (colour filled by sample site) and indicates collection site of the snail. Taxa represented on the plot are the top 20-ranked differentials in predicting whether a snail was from New Zealand or Europe and are annotated at the deepest available taxonomic level. Density plot below PCo1 shows snail sample density across PCo1 labelled by sample site. Density plots to the left of PCo2 show snail sample density across PCo2 labelled by sex (male or female), infection status (infected or uninfected), and reproductive mode (sexual or asexual).

To identify taxa that significantly predicted ecosystem clustering, where ecosystems are defined by the two regions of Europe and New Zealand, we used machine learning classification via a random forest model to classify snails as sampled from Europe or New Zealand based on microbiome composition. The model classified samples based on relative microbial abundance into European or New Zealand ecosystems with 99.6% accuracy. We corroborated these results using a reference frames approach and identified taxa that are associated with snails from Europe or New Zealand (52). There were eight phyla that were exclusively associated with snails from Europe and none that were exclusively associated with snails from New Zealand (Table 2; Fig. 1). However, there were more Firmicutes taxa associated with snails from New Zealand (9 taxa) than Europe (6 taxa) (Table 2; Table S1).

**Table 2.** Europe and New Zealand microbiome phyla predictors. We used a multinomial regression to predict whether snails were from Europe or New Zealand samples sites. Represented here is a summary of counts of phyla that were associated with snails from EU or NZ.

| **Phylum** | **Europe** | **New Zealand** |
| --- | --- | --- |
| Acidobacteria | 3 | 0 |
| Actinobacteria | 4 | 0 |
| Armatimonadetes | 2 | 0 |
| Bacteroidetes | 31 | 13 |
| Chlamydiae | 0 | 0 |
| Chloroflexi | 0 | 0 |
| Cyanobacteria | 4 | 3 |
| Deinococcus-Thermus | 1 | 0 |
| Epsilonbacteraeota | 1 | 0 |
| Firmicutes | 6 | 9 |
| Gemmatimonadetes | 0 | 0 |
| Lentisphaerae | 0 | 0 |
| Nitrospirae | 1 | 1 |
| Planctomycetes | 2 | 0 |
| Proteobacteria | 65 | 32 |
| Spirochaetes | 1 | 0 |
| Tenericutes | 1 | 0 |
| Verrucomicrobia | 1 | 2 |

To test the degree to which microbiota exhibit host population-specific distributions, we calculated the core microbiome of snails from Europe and New Zealand, defined at the tip agglomerated level. European snails had a core microbiome (found in >=90% of samples) consisting of 30 taxa while snails from New Zealand contained a core of one taxon (Fig 2A), *Arenimonas*, which was also a part of the European snail sore microbiome. Our results departed from this prediction of higher core microbiome specificity for sites within the New Zealand region by showing relatively consistent core microbiome composition, containing fewer taxa, across the New Zealand sample sites compared to European sample sites. We found that the core microbiome of European snails constituted, on average, 13.8% of their microbiome. By contrast, the core microbiome of New Zealand snails constituted 70.1% of their microbiome, a marked and statistically significant difference between native and invasive snails (Fig 2B; T-test; *p* < 0.01). Proteobacteria were the main constituent of the core microbiota in both snail populations, making up 100% (1/1) of the New Zealand snail core and 68.9% of the taxa in the European snail core (20/29).

**Figure 2.**
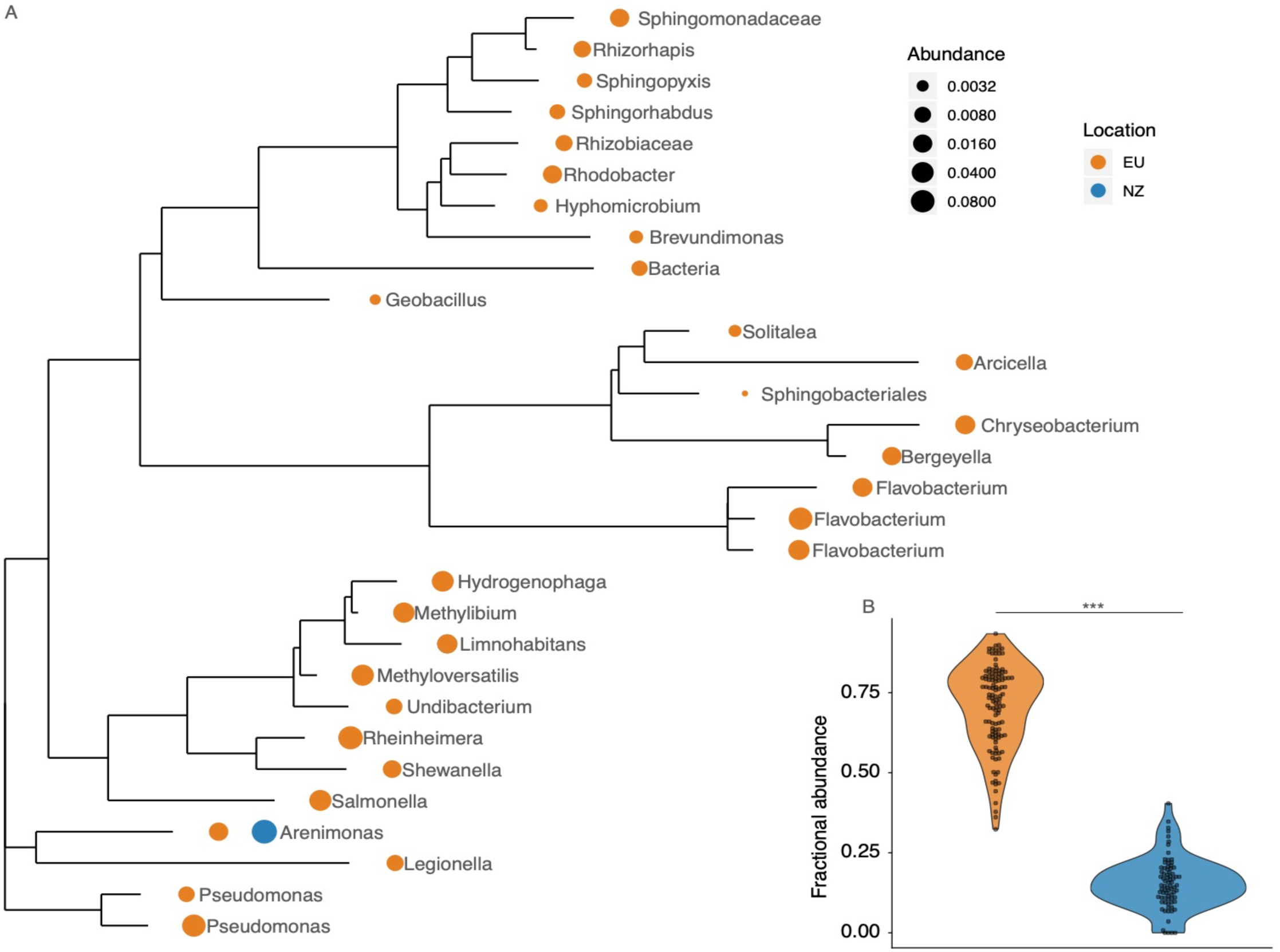
Core microbiota of European and New Zealand snails. (A) 16S rRNA phylogeny of core microbiota (>90% samples) of European (EU) and New Zealand (NZ) snails. Fractional abundance averaged across samples within the EU and NZ regions is represented by dot size.(B) Violin plot summarizing the proportion of amplicon reads mapped to core. Significance is denoted as *** = *p* < 0.001 (T-test).

After finding that the majority of predictive taxa were in higher relative abundance in European samples and most taxa from the core microbiome of European samples were not found in New Zealand, we next evaluated whether there were differences in species richness between European and New Zealand snails. We found that European snails (mean: 116 ASVs, s.e.: 2.51) had on average 3.52x more ASVs than New Zealand snails (mean: 32.9 ASVs, s.e.: 2.56) (Fig. 3; T-test; adj*-p* < 0.01; t = −23.1). There was significantly higher Chao1 and PD Whole Tree diversity amongst European snail microbiota, suggesting higher richness of rare species at relatively low abundance and higher phylogenetic diversity in European snail microbiota compared to New Zealand snail microbiota (Fig. 3; T-test; adj*-p* values < 0.01; Chao1 t = −25.2; PD t = −24.8). Shannon diversity was also significantly higher in European snail microbiota (Fig. 3; T-test; adj*-p* < 0.01; t = −4.52). Our subsequent prediction that New Zealand snail microbiota may have higher Pielou’s evenness than the European snails was upheld (Fig. 3; T-test; adj*-p* < 0.01; t = 22.4). To confirm that alpha diversity differences between microbiomes of snails from Europe and New Zealand were not due to uneven sampling (10 European sample sites vs. 3 New Zealand samples sites), we permuted alpha diversity tests on all combinations of three Europe sites vs. the three New Zealand sites (Fig. S3). Amongst the 600 combination tests, 98.3% remained significant (Table S2; T-test; adj*-p* < 0.05), where the non-significant findings were for Shannon diversity comparisons between European sites including West Holme and Melle compared to New Zealand sites. There were thus more species, more rare species, and higher phylogenetic richness in European snail microbiota, as well as higher species evenness in New Zealand snail microbiota.

**Fig 3.**
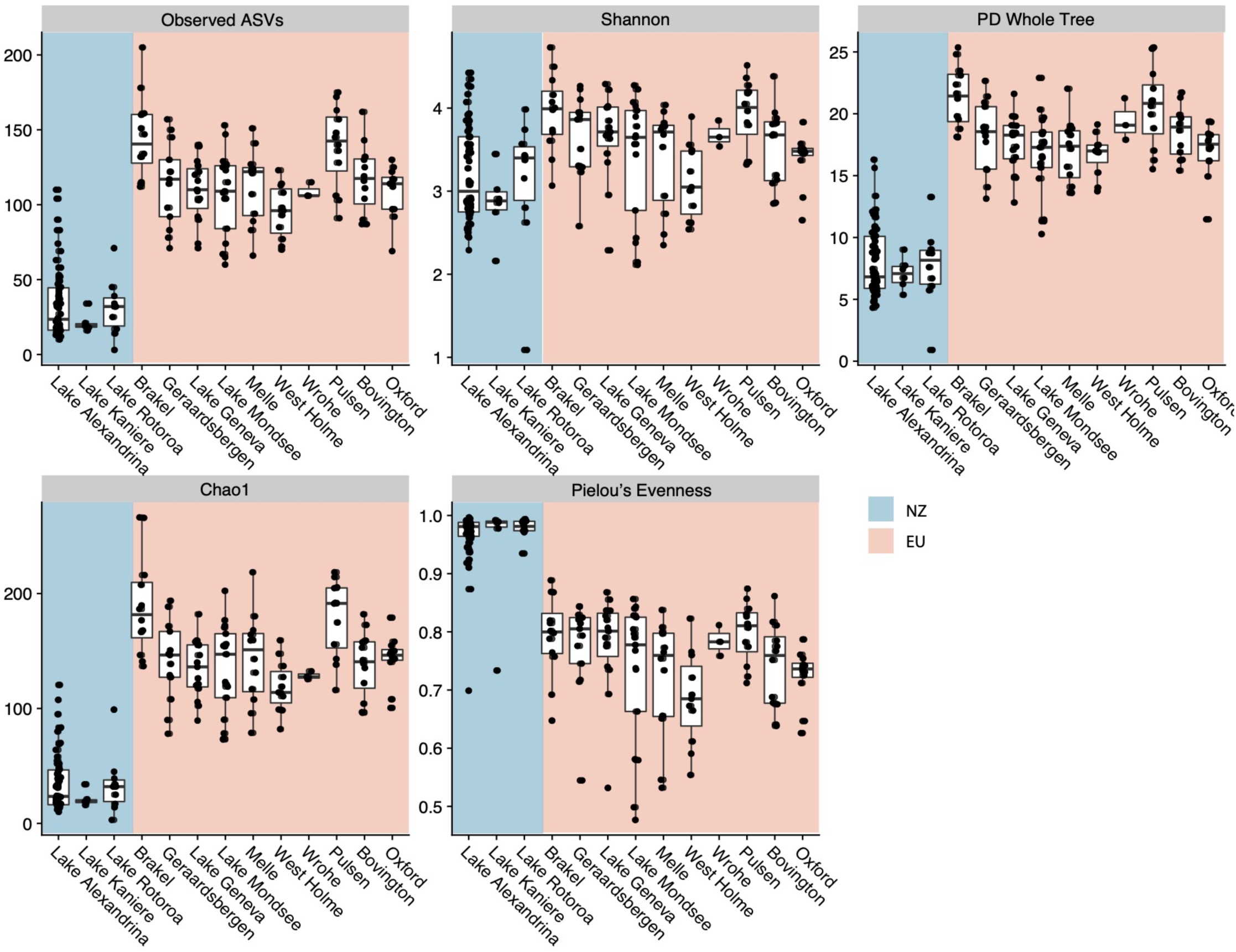
Alpha diversity measurements across sample sites. Figures are faceted by diversity metric. X-axis shows sampling site. Y-axis shows value of diversity metric. NZ = New Zealand samples and EU = European samples.

Our comparisons of snails from Lake Alexandrina show that sex (R^2^ = 0.147; adj-*p* = 0.001), reproductive mode (R^2^ = 0.0467; adj-*p* = 0.001), and infection status (R^2^ = 0.0330; adj-*p* = 0.005) all are associated with microbiota composition (Table 3; ADONIS; Permutations: 999). We controlled for these covariate effects and highlight taxa that significantly differed in abundance by sex and reproductive mode (Fig. S4; ANCOM; adj-*p* < 0.05). No taxa significantly differed in abundance based on infection status. We identified ten ASVs (all Xanthomonadaecae) that were significantly higher in abundance in asexual vs. sexual snails (Fig S4). We observed 50 ASVs (*Niabella* (n = 16), *Bacillus* (n = 15), and OM60 (n = 19)) that were significantly more abundant in male vs. female snails. There was significantly higher ASV richness in uninfected compared to infected snails (T-test; adj*-p* < 0.01; Fig S5). There were no significant differences in observed ASV richness when comparing reproductive mode or sex (Fig. S5).

**Table 3.** ADONIS testing effects of sex, ploidy, and infection status on microbiome UniFrac distances. Results from Analysis of Variance Using Distance Matrices (ADONIS) testing of the effects of sex, ploidy, and infection status on unweighted microbiome UniFrac distances between snails. All tests were performed on snails from Lake Alexandrina with 999 permutations.

| **Condition** | **Df** | **SS** | **MS** | **F Model** | **R2** | **P** |
| --- | --- | --- | --- | --- | --- | --- |
| **Sex** | 1 | 1.2899 | 1.28991 | 14.5946 | 0.14665 | 0.001 |
| **Reproductive mode** | 1 | 0.4105 | 0.41049 | 4.6444 | 0.04667 | 0.001 |
| **Infection** | 1 | 0.29 | 0.29 | 3.2813 | 0.03297 | 0.005 |
| **Residuals** | 77 | 6.8055 | 0.08838 | 0.77371 |  |  |
| **Total** | 80 | 8.7959 | 1 |  |  |  |

## DISCUSSION

There is a potential for loss of mutualistic microbes upon invasion due to subsampling and/or the reduced time for host-microbiota coevolution in the new range. Thus, we predicted that invasion of new habitat by a host organisms would reflected in changes to the composition and/or diversity of host-associated microbes. The native New Zealand and invasive European snails had distinct microbiotas. Whilst invasive snails carried the core taxa from New Zealand, they harboured more core taxa overall and higher phylogenetic diversity. These results suggest that invaders seem to readily acquire new symbionts, which may play a role in the success of this worldwide invader. Because numerous new microbes appear to be readily acquired by invasive snails, our results also suggest that the microbes associated with invasive lineages may be opportunistic and potentially less specialized. This pattern may be a consequence of the comparatively short time for host-microbe coevolution, relative to the long coevolutionary time between microbes and New Zealand snails, such that host-control mechanisms have not yet evolved to keep microbial diversity in check (19).

The core microbe from native New Zealand snails was found in European invasive snails. This result suggests that there could be selection to maintain certain symbiont associations in both ranges, which may be important to host biology. Because we were unable to obtain environmental samples due to insufficient amounts of DNA in water samples, and it remains unclear whether some core microbes are inherited, we cannot say whether core microbes found in European and New Zealand populations were present in both environments, or carried over from the latter. Nevertheless, European samples had 30 times more core taxa overall, and their most predictive taxa were absent or relatively infrequent in native snails. In other systems, it has been shown that invaders have lower microbial diversity and/or a subset of microbes relative to that present in native populations (15, 16). Invasive *P*. *antipodarum* populations feature low genotypic diversity relative to the native range (31, 54, 55) and mitochondrial haplotypes of invaders represent genetic subsets of native mitochondrial haplotype diversity (17). However, this genetic subsampling in invasive *P*. *antipodarum* populations is not reflected in their microbiota diversity.

Across both ranges, proteobacteria dominate among the core microbes. Proteobacteria and the majority of the other core microbial taxa observed here are consistent with Takacs-Vesbach et al. (26). The predominance of proteobacteria among the core microbes suggest a shared coevolutionary history between snails and microbial symbionts has driven the loss of non-essential partners, and ultimately streamlined the microbiome down to specialized taxa (19). A similar phenomenon occurs in a number of insect systems, whereby the host harbours few core microbes that are key players in organismal function (56, 57). Conversely, other insect taxa are host to relatively diverse core microbial populations (58).

It is well established that antagonistic coevolutionary interactions such as between hosts and pathogens can drive rapid evolution and diversification (59, 60). In the case of mutualisms, theory has predicted relatively low rates of evolution compared to antagonistic relationships (61, 62), although empirical evidence in support of this conjecture remains mixed (63). For example, mutualisms can drive diversification, such as that observed in plant-animal mutualisms (64). Even in the absence of host evolution, beneficial resident microbes can evolve within days (65). Furthermore, competitive interactions between microbes that overlap in resource use can also drive diversification (20).

Coevolution between *P*. *antipodarum* and *A*. *winterbourni* is well documented (27, 29, 30). Reciprocal antagonistic selection between *P*. *antipodarum* and *A*. *winterbourni* is expected to be strong because infected snails are completely sterilized (66), and parasites die if they encounter resistant individuals (67). In other systems, the potential for mutualist microbes to be involved in host defence and even host-parasite coevolution has been demonstrated (68, 69). However, the extent to which coevolution operates in diverse communities that feature both mutualists and antagonists is unclear. Our results do not suggest that the microbiota is a dominant player in this host-parasite coevolutionary relationship: some taxa differed only marginally, but not statistically significantly, in relative abundance between infected and uninfected snails, although we did observe significantly higher ASV richness in uninfected snails compared to infected snails.

In other systems, it does seem that microbial diversity could confer some degree of parasite resistance (70), perhaps analogous to the host genetic diversity conferred by outcrossing in *P*. *antipodarum* (71). However, asexual *P*. *antipodarum* are more successful invaders strongly, which suggests that this diversity cannot outweigh the costs of being sexual. Here, we found that reproductive mode was associated with microbiome composition and ten ASVs that were significantly more abundant in asexuals than sexuals. Takacs-Vesbach et al. (26) observed significant differences in microbiome composition between sexuals and asexuals and among lake populations of *P*. *antipodarum*. Because the mechanism(s) leading to transitions from diploid sexuality to polyploid asexuality in *P*. *antipodarum* are still unclear, future work could focus on the possible links between microbes and reproductive mode and/or polyploidization. For example, *Wolbachia*-mediated transitions to asexuality are well characterized in arthropods (72). In regard to potential effects of ploidy level, Takacs-Vesbach et al. (26) did not observe significant differences in in microbiome composition between asexual triploids and tetraploids, and we were unable to obtain sufficient sampling of tetraploids to perform ploidy level comparisons. Furthermore, besides a handful of studies (73, 74), very little is known about potential links between host ploidy level and microbiome composition, especially in animals

While it is established that symbiotic microbes have important effects on host fitness, the relationship between microbes and molluscan invaders has received little attention. Here, we find that the process of invasion is associated with some carryover of a core microbiota, but mostly involves the formation of numerous new symbiotic relationships, suggesting that invasive *P*. *antipodarum* are able to take advantage of readily available microbes. Despite some impact of biological trait variation on the microbiota in native snail populations, there is a consistent and strong pattern suggesting that host-microbiota coevolution (or host evolution alone (19)) streamlines the microbiota of native snails to essential members. Future research will need to establish the fitness impacts of those new microbes on invaders, and whether the invading snails have adapted to accommodate them, which could be accomplished by reciprocal microbial transplant experiment in which germ-free snails are exposed to snail-associated microbes from different environments. Notwithstanding this possibility, we show that the potential for invasion to alter the microbiome is clear. Future research seeking to control invasive species should focus on the new associations being formed with microbes which might be contributing substantially to invader success.

## DATA AVAILABILITY

Raw sequencing data will be made publicly available on NCBI Sequence Read Archive and accession numbers will be added upon article acceptance.

## ACKNOWLEDGEMENTS

We thank Katelyn Larkin, Dunja Lamatsch, Tanja Schwander, Kaitlin Hatcher, Bennett Brown, Kyle McElroy, and Curtis Bankers for assistance with snail field collections. We are grateful to the King Lab for feedback on the results, especially Kieran Bates for manuscript comments. Sequencing was conducted at the W.M. Keck Center for Comparative Functional Genomics (University of Illinois). Flow cytometry data were obtained at the Flow Cytometry Facility, a Carver College of Medicine/Holden Comprehensive Cancer Center core research facility at the University of Iowa. This work was supported by the University of Iowa T. Anne Cleary International Dissertation Research Fellowship (LB), the University of Iowa Center for Global and Regional Environmental Research Travel Award (LB), a BBSRC Doctoral Training Grant (BB/M011224/1, DD), and by a British Ecological Society Grant (KCK).

## SUPPLEMENTARY FIGURE AND TABLE CAPTIONS

**Fig. S1. Map of sampling sites**. (A) Locations of our ten European sampling sites. (B) Locations of our three New Zealand sampling sites. Country names and boundaries are in yellow. Sampling sites are represented by red dots and the location names are in white. Zoomed out inlayed maps show the broader regions with the sampled regions highlighted in the red boxes. Maps were made using Google Earth Pro v. 7.3.2.5776.

**Fig. S2. Snail microbiota ecosystem clustering**. Biplot PCoA on weighted UniFrac dissimilarity distances between snail microbiota profiles across European and New Zealand sampling sites. Each point represents a single snail, and points are colour-coded by whether a point is a microbial taxon or snail and indicates snail collection site. Taxa represented on the plot are the top 20-ranked differentials in predicting whether a snail is from New Zealand or Europe and are annotated at the deepest available taxonomic level. Density plot below PCo1 shows snail sample density across PCo1 labelled by sampling site. Density plots to the left of PCo2 show snail sample density across PCo2 labelled by sex (male or female), infection status (infected or uninfected), and reproductive mode (sexual or asexual).

**Fig. S3. Combination PCoAs on unweighted UniFrac dissimilarity distances between snail microbiota profiles across European and New Zealand sample sites**. Each point represents a single snail, and points are colour-coded by collection site of snail and shaped by whether a snail is from Europe or New Zealand. Include all combinations of snails from three European sites compared to snails from all three New Zealand sites.

**Fig. S4. ASVs that significantly differed in abundance in snail microbiota based on reproductive mode or sex**. Points represent significantly differentially abundant taxa and are plotted by log2fold change in abundance (ANCOM; adj-p < 0.01). Comparisons are asexual/sexual and male/female. There were no significantly differentially abundant taxa between infected and uninfected snails. Figure is labelled by phyla and deepest available taxonomy, where G = genus and F = family. Snails are only from Lake Alexandrina, NZ.

**Fig. S5. Observed ASVs faceted by infection status, reproductive mode, and sex**. Each point represents a snail sample. Comparing observed ASVs, as a measure of species richness, across snail metadata variables of infection status, reproductive mode, and sex. Data from Lake Alexandrina sails, the only collection site where all metadata variables were tested (T-test; *** = adj-p < 0.01).

**Table S1**. Complete list of differential taxa.

**Table S2. Combination alpha diversity comparisons**. Combinations include all combinations of snails from three European sites compared to snails from all three New Zealand sites (T-tests; FDR-corrected p-values).

## Notes

### Competing Interest Statement

The authors have declared no competing interest.

