## Supplementary Methods for "Invasive freshwater snails form novel microbial relationships"

**SUPPLEMENTARY MATERIALS AND METHODS**

**Sample Collection, Dissection, and Reproductive Mode Determination**

We used nets to collect adult *Potamopyrgus antipodarum* during warm seasons from shallow (lake depth < 1 m) rocks and vegetation from three New Zealand collection sites in January 2015 (Table 1; Fig. S1). After transport to the University of Iowa, these snails were maintained in separate identical 15 L tanks, one tank per source population, in a 16° C room with a light:dark cycle of 16:8 hours for less than one month before dissection. Snails were fed dried *Spirulina* cyanobacteria *ad libitum* (1). Invasive *P. antipodarum* from five countries in Europe (Table 1; Fig. S1) were collected in the same manner in spring 2016, transported to the University of Oxford, and maintained under the same conditions and for the same amount of time as were the New Zealand snails.

All snails used in this study were adults and were sexed and shells were removed prior to DNA extraction. Snails were assessed for *A. winterbourni* infection based on the presence of metacercarial cysts via dissection (2) (Table 1). All metacercariae were removed from infected snails using a micropipette. Snails containing non- *A. winterbourni* infections were excluded from the study. Dissected New Zealand snails were separated into two tubes, one tube containing one half of a head, which was split between the two tentacles, and the other tube containing the snail body and other head half. All samples were snap frozen in liquid nitrogen immediately following dissection and then stored at -80°C. The first head halves were used for flow cytometry (3, 4) to determine ploidy status as a proxy for reproductive mode (diploids are sexual and polyploids are asexual). The second head half and the whole body were used together for DNA extraction. Because it is already established that the European invasive lineages are virtually all polyploid asexuals (5, 6), we did not perform flow cytometry on these samples. Thus, the whole head and body from these snails were placed into single tubes and treated otherwise similarly to New Zealand samples.

**DNA Extraction and Library Preparation**

Whole-snail tissue (excluding the shell and parasite metacercariae) was lysed using a mortar and pestle in the Qiagen DNeasy Plant Mini Kit (QIAGEN Inc.) lysis buffer and then extracted following manufacturer protocol, but eluting DNA in 40 µl 100:1 TE buffer. The Plant kit was used as it better handles the polysaccharides present in snail mucus, compared to other DNA extraction kits. Following extraction, we analysed 1.5 µl of each sample on a Nanodrop® 1000 (Thermo Fisher Scientific) to determine the concentration, amount (ng), and quality of each DNA extraction. Samples with a 260/280 ratio > 1.6 and containing > 20 ng of total DNA were used in further analyses. We ran 3 µL of the eluted DNA on a 1% agarose gel for each sample that achieved the above quality criteria and photographed each gel using a FOTODYNE imaging system (FOTODYNE Inc.). Samples that reached our quality criteria and produced clear bands on a gel were shipped to the W.M. Keck Center for Comparative Functional Genomics (University of Illinois at Urbana-Champaign) for sequencing. DNA extractions were stored at -80°C until library preparation.

The 16S rRNA V4 region was amplified from the *P. antipodarum* microbiome gDNA using the 515F Golay-barcoded primers and 806R primers (7, 8) listed on the Earth Microbiome Project (EMP) 16S protocol site (http://www.earthmicrobiome.org/emp-standard-protocols/16s/). Samples were prepared in accordance with the standard EMP 16S rRNA protocol (9). Our 25 $\mu$l polymerase chain reactions (PCR) contained 10 $\mu$l Platinum Hot Start MM (2X) (Thermofisher Scientific), 11 $\mu$l nuclease-free water, 1 $\mu$l of each forward and reverse primer (0.2 $\mu$M final concentrations), and 2 $\mu$l gDNA template. No-template controls (NTCs) contained nuclease-free water instead of gDNA. Reactions were held at 94°C for 3 min to denature the DNA, and amplification took place for 35 cycles at 94°C for 45 sec, 50°C for 60 sec and, 72°C for 90 sec. The cycles were followed by a hold at 72°C for 10 min. Amplicons were visualized on a 1.5% agarose gel. gDNA was quantified using the Qubit 2.0 fluorometer (Thermofisher Scientific,) and amplicons were pooled at equimolar ratios (~240 ng per sample). The combined amplicon pool was then cleaned using the Qiagen PCR Purification Kit (QIAGEN Inc.). The multiplexed library was quality checked and sequenced with the MiSeq 2x250 bp PE v2 protocol at the W.M. Keck Center for Comparative Functional Genomics (University of Illinois).

**Computational and Statistical Analyses**

We removed PhiX sequences from the sequencing libraries using Bowtie2 (10) by mapping reads against an index built from a PhiX genome (obtained from: support.illumina.com/sequencing/sequencing_software/igenome.html). We demultiplexed paired-end fastq files, which were then processed in R (3.4.0) using DADA2 as previously described (11). In short, this process included filtering and trimming, error rate estimation, de-replication of reads into unique sequences, and amplicon variant inference. We used the suggested filter and trimming parameters (11); truncated Q score (truncQ) was 2; forward and reverse reads were truncated at base pairs 240 and 160, respectively; maximum expected error (maxEE) for forward and reverse reads was 2; reads with more than zero Ns (maxN) were discarded; and PhiX reads were removed. We then merged paired-end reads, constructed an amplicon sequence variant (ASV) table, which is a sample-by-sequence abundance matrix, and removed chimeras. We also used the native implementation of the DADA2 Ribosomal Database Project (RDP) naïve Bayesian classifier (12) trained against the GreenGenes 13.8 release reference fasta (https://zenodo.org/record/158955#.WQsM81Pyu2w) to classify ASVs taxonomically.

We used the phyloseq v. 1.16.2 estimate_richness and vegan's pd function to calculate alpha diversity measurements of observed ASVs, Shannon's index, PD Whole Tree, Pielou's evenness, and Chao1 (13). Phyloseq was also used to perform ordinations using PCoA on unweighted and weighted UniFrac distance scores (14). We used a reference frames approach in the Songbird package to calculate taxa differentials (15). We also used the R packages ggplot2 for data visualization and figure generation (2.0.0) (16), Rcpp for C++ parallelization in R (17), optparse (1.3.2.) to parse command line options, stats (3.2.3) to run statistical analyses, and data.table (1.9.6) to handle data frames.

We controlled for effects of snail sex, reproductive mode, and infection status (Table 1) when testing for geographic associations with microbiota by only conducting analyses on female uninfected asexual (polyploid) snails, allowing us to directly compare native and invasive snails without the confounding factors of sex, reproductive mode, and infection (comparisons of sex, reproductive mode, and infection status are described below). We did this because all snails from Europe were female uninfected asexual (polyploid) snails. For alpha diversity analyses, we rarefied samples to 10,000 ASVs per sample and discarded two samples that had fewer reads than this threshold. To test covariate effects on microbiota alpha diversity, we used Welch's Two Sample t-tests and adjusted *p*-values ("adj-*p*") with a Bonferroni correction for multiple tests. We conducted beta diversity analyses on all ASVs after removing singletons. We normalized ASV counts by adding one and then $\log_e$-transforming ASV counts (11). To calculate beta diversity we first built distance matrices based on the unweighted and weighted UniFrac scores of each sample (14) and then performed PCoA on the distance matrices.

To evaluate the effect of geographic location (sample site) on microbiota beta diversity, we used Analysis of Similarity (ANOSIM) tests. To avoid the possible confounder that all European snails were female uninfected asexual (polyploid) snails and many New Zealand snails were sexual (diploid), infected, and/or male, we only performed the ANOSIM comparing Europe and New Zealand on female uninfected asexual snails. ANOSIM tests were conducted with 999 permutations, and ANOSIM R-statistics ($R^2$) and exact *p*-values are reported in the results.

For our analysis of population specificity, we focused on the core microbiome of snails within locations, defined as taxa found in at least 90% of samples from a given location. We calculated the proportion of shared core microbiome taxa among all sampling locations and also compared all European samples vs. all New Zealand samples. We used a t-test to compare the proportion of reads that mapped as core microbiome taxa between the combined Europe and combined New Zealand sample groupings.

We performed a Permutational Multivariate Analysis of Variance Using Distance Matrices analysis (ADONIS) to test the effects of reproductive mode, sex, and infection status on microbiota beta diversity. The ADONIS analysis was limited to the Lake Alexandrina (New Zealand) sample site as it was the only site for which we were able to obtain all conditions (sexual, asexual, male, female, infected, and uninfected). ADONIS tests were conducted with 999 permutations.

We employed the Analysis of Composition of Microbes (ANCOM) algorithm to conduct a differential count analysis (18). After observing that geography, sex, reproductive mode, and infection status significantly affected beta diversity (see Results), we corrected for these associations in our differential count comparisons by sex, reproductive mode, and infection status. This set of analyses was limited to the Lake Alexandrina (New Zealand) sample site as it was the only site for which we were able to obtain all conditions (sexual, asexual, male, female, infected, and uninfected). We tested the effect of sex in uninfected male vs. uninfected female sexual snails, tested the effect of reproductive mode on uninfected female sexual vs. uninfected female asexual snails, and tested the effect of infection status on asexual female trematode-infected vs. asexual female uninfected snails. For our machine learning approach to model microbiome classification by geography, we again used female uninfected asexual snails and ASVs agglomerated phylogenetically using default settings (h = 0.2) in phyloseq's tax_glom function. We performed machine learning classification training with a random forest model using the caret package (v6.0-81). We used a training test split of 80:20 and fit a random forest classifier over the tuning parameter of snail origin (New Zealand or Europe).
