## Supplementary material for "Invasive freshwater snails form novel microbial relationships": Fig. S1

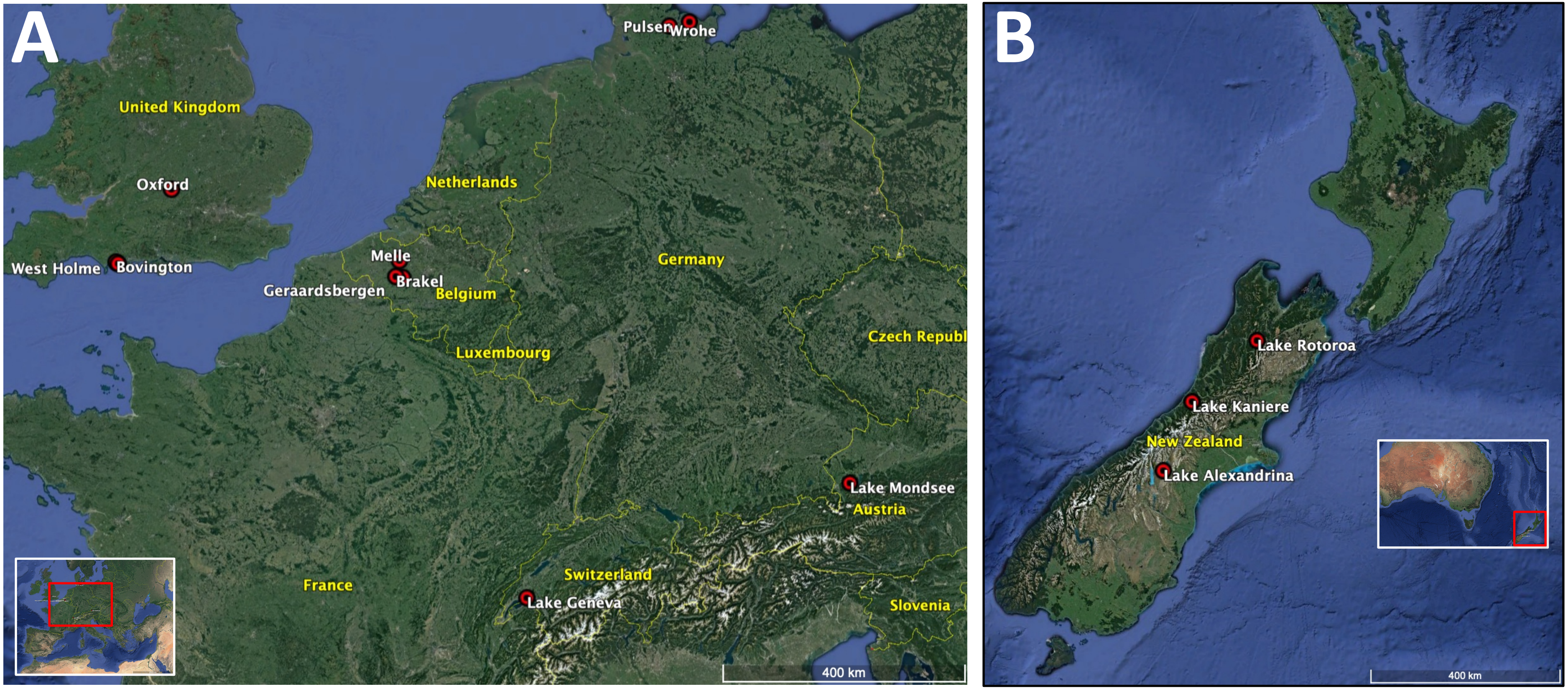


**Fig. S1. Map of sample sites.** (A) Locations of our ten European samples sites. (B) Locations of our three New Zealand sample sites. Country names and boundaries are in yellow. Sample sites are represented by red dots and the location names are in white. Zoomed out inlayed maps show the broader regions with the sampled regions highlighted in the red boxes. Maps were made using Google Earth Pro v. 7.3.2.5776.
