## Supplementary material for "Invasive freshwater snails form novel microbial relationships": Fig. S2

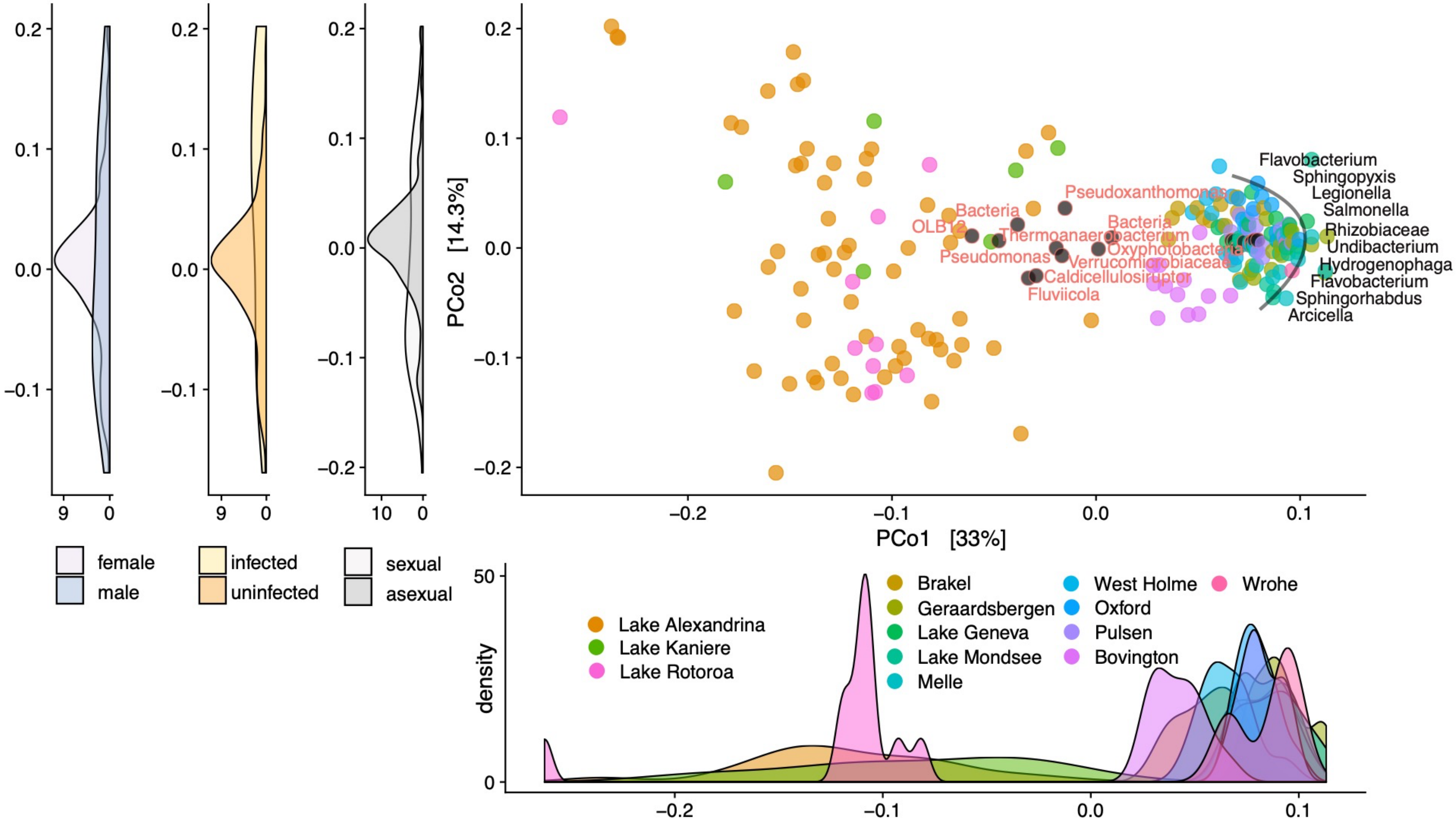


**Fig. S2. Snail microbiota ecosystem clustering.** Biplot PCoA on weighted UniFrac dissimilarity distances between snail microbiota profiles across European and New Zealand sample sites. Each point represents a single snail, and points are colour-coded by whether a point is a microbial taxon or snail and indicates collection site of snail. Taxa represented on the plot are the top 20-ranked differentials in predicting whether a snail is from New Zealand or Europe and are annotated at the deepest available taxonomic level. Density plot below PCo1 shows snail sample density across PCo1 labelled by sample site. Density plots to the left of PCo2 show snail sample density across PCo2 labelled by sex (male or female), infection status (infected or uninfected), and reproductive mode (sexual or asexual).
