## Supplementary figures and images for "Invasive freshwater snails form novel microbial relationships"

### Fig. S3

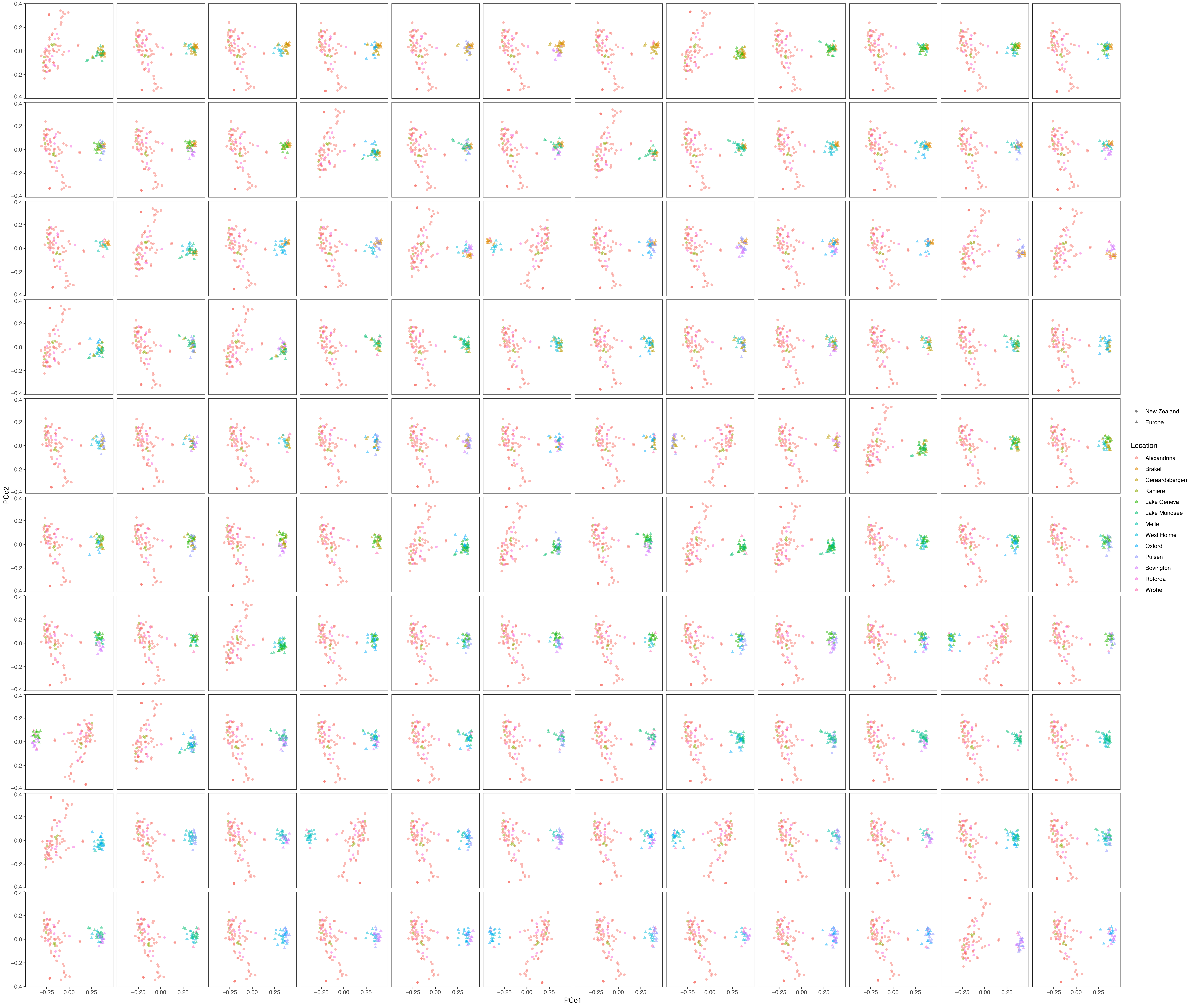
