## Supplementary material for "Invasive freshwater snails form novel microbial relationships": Fig. S4

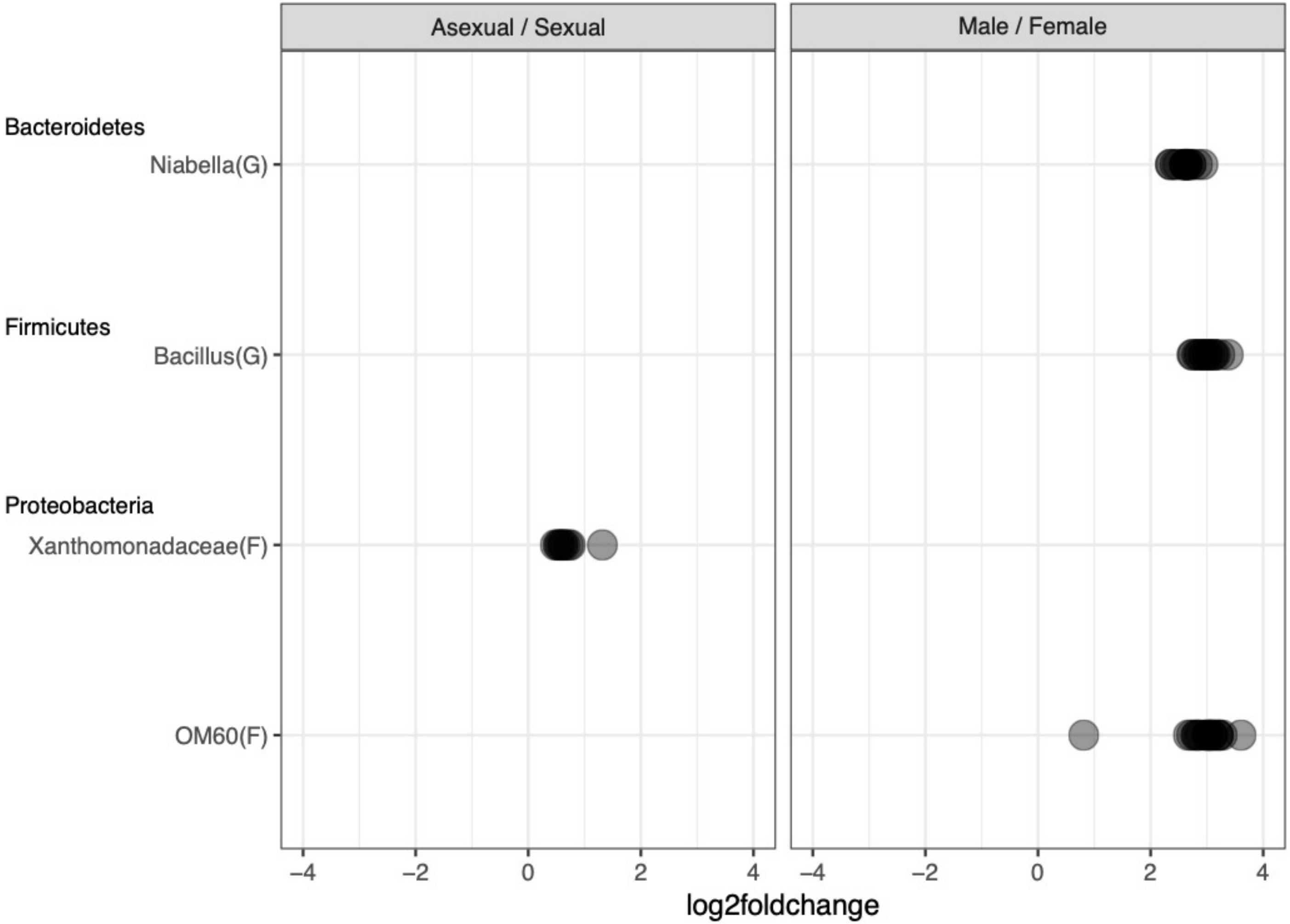


**Fig. S3. ASVs that significantly differed in abundance in snail microbiota based on reproductive mode or sex.** Points represent significantly differentially abundant taxa and are plotted by $\log_2$fold change in abundance (ANCOM; adj-*p* < 0.01). Comparisons are asexual/sexual and male/female. There were no significantly differentially abundant taxa between infected and uninfected snails. Figure is labelled by phyla and deepest available taxonomy, where G = genus and F = family. Snails are only from NZ.
