## Supplementary material for "Invasive freshwater snails form novel microbial relationships": Fig. S5

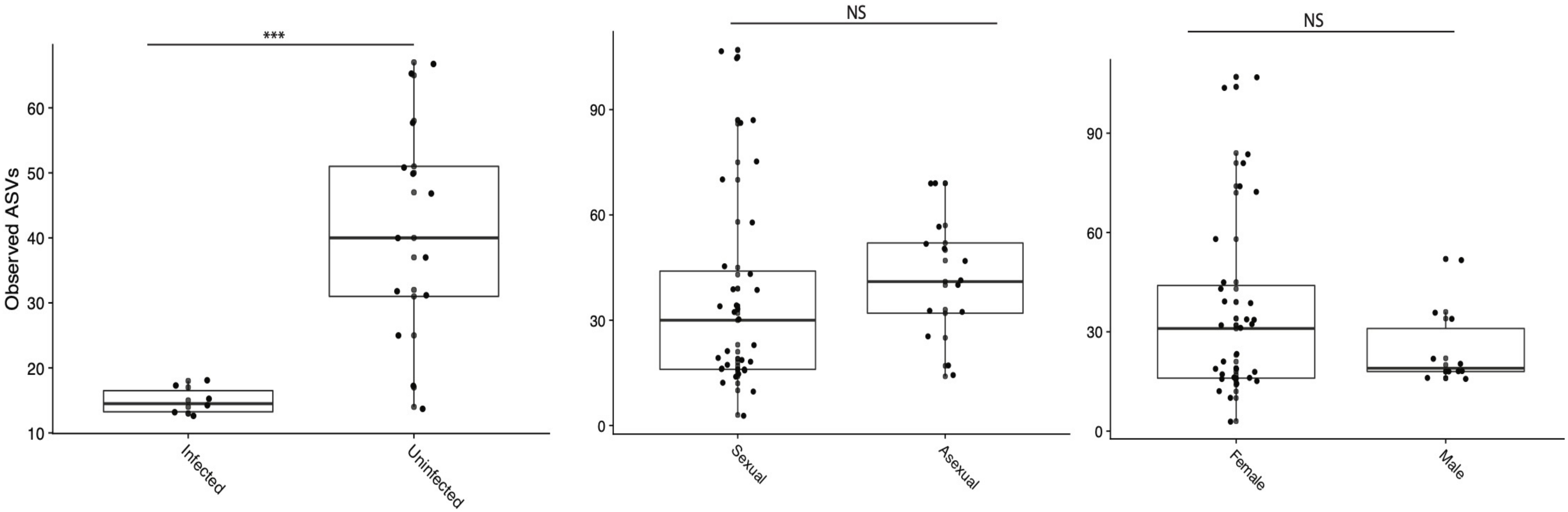


**Fig. S4. Observed ASVs faceted by infection status, reproductive mode, and sex**. Each point represents a snail sample. Comparing observed ASVs across snail metadata variables of infection status, reproductive mode, and sex. Data is of snails from Lake Alexandrina, the only location where all metadata variables were tested (T-test; *** = adj-$p < 0.01$).
